# Multi-parametric MRI can detect enhanced myelination in the *Gli1^-/-^* mouse brain

**DOI:** 10.1101/2023.11.20.567957

**Authors:** Choong H. Lee, Mara Holloman, James L. Salzer, Jiangyang Zhang

**Author notes:** Corresponding to: Jiangyang Zhang, Ph.D., Center for Biomedical Imaging, Department of Radiology, New York University School of Medicine, 660 First Ave, New York, NY 10016, USA.

## Abstract

This study investigated the potential of combining multiple MR parameters to enhance the characterization of myelin in the mouse brain. We collected *ex vivo* multi-parametric MR data at 7 Tesla from control and *Gli1*^−/−^ mice; the latter exhibit enhanced myelination at postnatal day 10 (P10) in the corpus callosum and cortex. The MR data included relaxivity, magnetization transfer, and diffusion measurements, each targeting distinct myelin properties. This analysis was followed by and compared to myelin basic protein (MBP) staining of the same samples. Although a majority of the MR parameters included in this study showed significant differences in the corpus callosum between the control and *Gli1*^−/−^ mice, only T_2_, T_1_/T_2,_ and radial diffusivity (RD) demonstrated a significant correlation with MBP values. Based on data from the corpus callosum, partial least square regression suggested that combining T_2_, T_1_/T_2_, and inhomogeneous magnetization transfer ratio could explain approximately 80% of the variance in the MBP values. Myelin predictions based on these three parameters yielded stronger correlations with the MBP values in the P10 mouse brain corpus callosum than any single MR parameter. In the motor cortex, combining T_2_, T_1_/T_2,_ and radial kurtosis could explain over 90% of the variance in the MBP values at P10. This study demonstrates the utility of multi-parametric MRI in improving the detection of myelin changes in the mouse brain.

## 1. Introduction

Myelin is a multi-layered structure with lipid-rich cell membrane that is generated by the oligodendrocytes to insulate and support axons to facilitate rapid and efficient conduction of neuronal signals (Chang et al., 2016). The main chemical composition of myelin is lipid (∼80 % of its dry mass) (O’Brien and Sampson, 1965), with proteins comprising only ∼20% of its dry mass, including proteolipid protein (PLP) and myelin basic protein (MBP). Myelin pathology is commonly found in neurological diseases, either with myelin as a direct target, such as in multiple sclerosis (MS) (Chang et al., 2016), leukodystrophy (van der Knaap and Bugiani, 2017), or indirectly as in Alzheimer’s disease (Chen et al., 2021; Depp et al., 2023). Thus, myelin is a central therapeutic target in primary demyelinating disorders. It is therefore important to develop methods for imaging myelin in order to detect myelin injury and monitor its repair, especially non-invasive imaging techniques.

MRI provides a myriad of tissue contrasts for the detection of myelin content and injury, including tissue relaxivity (e.g., T_1_, T_2_) (Glasser et al., 2014; Laule et al., 2008; Laule et al., 2006; Laule et al., 2011; MacKay and Laule, 2016; Miot-Noirault et al., 1997), susceptibility (Duyn, 2013; Lee et al., 2012; Shin et al., 2021), magnetization transfer (MT) (Henkelman et al., 2001; Rademacher et al., 1999; Sled, 2018; Vavasour et al., 2011; Xydis et al., 2006), and diffusion (Guglielmetti et al., 2016; Harkins and Does, 2016; Rahmanzadeh et al., 2021). T_1_ and T_2_-MRI has long been used to survey myelin pathology in the brain (Glasser et al., 2014; Mottershead et al., 2003), and the ratio between T_1_-weighted and T_2_-weighted MRI signals has been used as a measure of cortical myelin in the human brain (Glasser et al., 2014). Furthermore, based on the distinct relaxivity properties of water between myelin sheath, the fraction of the myelin water compartment can be estimated, which provides a more direct and sensitive marker for myelin (MacKay et al., 1994; MacKay and Laule, 2016).

MT-MRI measures the transfer of magnetization between protons bounded to macromolecules (e.g. lipids and proteins) and free water protons (Henkelman et al., 2001; Wolff and Balaban, 1989). In a typical MT-MRI scan, a train of off-resonance radiofrequency pulses is applied to saturate the bounded protons, and some of the saturation is subsequently transferred to nearby free protons, effectively reducing the free protons’ signal. MT MRI can be particularly useful in studying tissues with abundant macromolecules, including myelin in the CNS (Hakkarainen et al., 2016; Schmierer et al., 2004; Vavasour et al., 2011), and quantitative MT has been developed to characterize tissue MT properties (Ramani et al., 2002; Sled and Pike, 2001; Yarnykh and Yuan, 2004). Recently, inhomogeneous MT (ihMT), a variant of MT-MRI, presented its potential as a myelin marker with enhanced sensitivity to myelin lipid (Varma et al., 2015a; Varma et al., 2015b). Duhamel et al. showed a strong correlation between ihMT signals and the myelin content of several mouse brain structures as measured using fluorescence signals in the *PLP1-GFP* mouse brain, where the green fluorescent protein (GFP) is expressed under the transcriptional control of the myelin PLP regulatory sequences (Duhamel et al., 2019; Spassky et al., 2001). We have also demonstrated that ihMT-MRI can detect residual non-compact myelin in the hypomyelinated shiverer mouse brain better than conventional MT-MRI (Lee et al., 2022).

The distinct microstructural organization of myelin e.g., the multilamellar wrapping of the myelin sheath around an axon, makes it a restrictive barrier to water molecule diffusion, which can be detected by diffusion MRI. Early reports suggested mean diffusivity (MD) (Schmierer et al., 2008; Thiessen et al., 2013; Yano et al., 2018) and fractional anisotropy (FA) (Schmierer et al., 2008; Thiessen et al., 2013; Wendel et al., 2018; Yano et al., 2018), two metrics derived from diffusion tensor imaging (DTI) (Basser et al., 1994; Mori and Zhang, 2006), might be sensitive to myelin. Later, radial diffusivity (RD), which measures the extent of water diffusion perpendicular to the axon, was demonstrated by Song et al. to be sensitive to demyelination (Song et al., 2005). Diffusion kurtosis, which characterizes non-Gaussian diffusion property (Jensen et al., 2005), has also been used to study myelin diseases (Guglielmetti et al., 2016; Kelm et al., 2016; Praet et al., 2018). There has been a strong interest in using biophysical model-based analysis to extract the properties of specific cellular compartments (e.g., neurites and soma) from diffusion MRI signals (Fieremans et al., 2011; Hansen et al., 2017). It has been shown that results from biophysical models can be combined with information from other contrast mechanisms to provide meaningful insights into myelin integrity and potential links to histology-based markers (e.g., g-ratio) (Jung et al., 2018; Stikov et al., 2015; Stikov et al., 2011).

The specificity of the above-mentioned and other MRI myelin markers, however, remain limited as demonstrated by recent meta-analyses (Lazari and Lipp, 2021; Mancini et al., 2020), mainly due to the indirect nature of MRI. This hinders the use of MRI to evaluate myelin change with precision (Patel et al., 2020). Previous studies have suggested that combining multiple MRI contrast mechanisms can benefit the characterization of myelin (Mangeat et al., 2015; Stikov et al., 2015; Stikov et al., 2011). In this study, we investigated whether combining multiple MR parameters, each targeting distinct aspect of myelin, can enhance the precision of MRI-based myelin mapping in the brain. We acquired multi-parametric MRI data from the brains of Gli1 knockout (*Gli1^-/-^*) and control mice. Loss of Gli1 promotes the differentiation of endogenous neural stem cells into oligodendrocytes thereby resulting in accelerated myelination of the developing forebrain compared to controls (Samanta et al., 2015). We first examined the ability of MR parameters to detect the difference in myelin between *Gli1^-/-^* and control brains using MBP-stained histology as the ground truth. We then used partial least square regression (PSLR) to identify a subset of MR parameters that were critical for myelin estimation and combinations of these selected MR parameters for precise myelin mapping in this model.

## 2. Methods

### 2.1. Animals

The animal protocol has been approved by the institutional animal care and use committee at New York University Grossman School of Medicine. We crossed *Gli1^+/-^* heterozygous (het) mice i.e., *Gli1^CreERT2^*and *Gli1^nLacZ^* to generate *Gli1^CreERT2/nLacZ^*i.e., *Gli1* null (*Gli1^-/-^*) mice; *Gli^+/-^*hets served as controls. Postnatal day 10 (P10) het and null mice were perfusion-fixed with 4% paraformaldehyde (n = 5 in each group). We chose the P10 time point as *Gli1^-/-^* mice are substantially myelinated at this time in contrast to control mice, which exhibit limited myelination (Samanta et al., 2015). In this study, we chose *ex vivo* MRI due to lengthy acquisition of multiple MRI parameters and the lack of fully developed teeth and ear canals in the P10 mouse for reliable motion constraints. Dissected mouse brains were placed in 10 ml syringe filled with Fomblin (perfluoropolyether; Ausimont USA Inc.), which has no MRI visible proton, to match tissue susceptibility and prevent dehydration.

### 2.2. MRI Acquisition

All samples were imaged at room temperature using a 7 Tesla MRI system with a quadrature transmit volume coil (70 mm diameter) and a 4-channel receive-only phased array cryogenic probe (Bruker Biospin, Billerica, MA, USA).

For T_1_ and T_2_ MRI, we used the RAREVTR (Rapid Acquisition with Relaxation Enhancement and Variable Repetition time (TR) (Hennig et al., 1986) and MSME (Multi-Slice Multi-Echo) sequence with the parameters in **Table 1**. For MT-MRI, we used an offset frequency of 5 kHz for conventional MT-MRI and 10 kHz for ihMT-MRI following previous reports by Prevost et al. (7). Co-registered diffusion tensor data were acquired using a diffusion-weighted echo-planar imaging (DW-EPI) sequence. T_1_/T_2_, MT/ihMT, DW-EPI images were acquired using the scan parameters shown in **Table 1**.

**Table 1:** MR imaging parameters used in this study. Abbreviations are: AT: acquisition time; NEX: number of excitation; TE: echo time; TR: repetition time.

| A | TE (ms) |  | TR (ms) |  | Resolution (mm<sup>3</sup>) |  | NEX/AT |
| --- | --- | --- | --- | --- | --- | --- | --- |
| T1-weighted RARE images | 6.3 |  | 868, 1096, 1365, 1694, 2115, 2702, 3682, 7500 |  | 0.1 x 0.1 x 0.7 |  | 2/15 min |
| T2-weighted MSME images | 7, 14, 21, 28, 34, 41, 48, 55, 62, 69, 76, 83, 89, 96, 103, 110, 117, 124, 131, 138, 144, 151, 158, 165, 172 |  | 4000 |  | 0.1 x 0.1 x 0.7 |  | 2/40 min |

| B | TE (ms) | TR (ms) | Offset Freq (kHz) | Echo Train Length | Saturation Pulse | Pulsewidth/Bandwidth/Repetition time/# of pulses | Resolution (mm<sup>3</sup>) | NEX/AT |
| --- | --- | --- | --- | --- | --- | --- | --- | --- |
| MT | 5.5 | 4000 | 5 | 6 | Gaussian | 1.83 ms/1.5 kHz/2.62ms/200 | 0.1 x 0.1 x 0.7 mm | 4/22 min |
| ihMT | 5.5 | 4000 | 10 | 6 | Gaussian | 1.83 ms/1.5 kHz/2.62ms/200 | 0.1 x 0.1 x 0.7 mm | 4/1hr 14 min |

| C | TE (ms) | TR (ms) | $\Delta/\delta$ (ms) | A0 images | Directions | B values (ss/mm<sup>2</sup>) | Resolution (mm<sup>3</sup>) | NEX/AT |
| --- | --- | --- | --- | --- | --- | --- | --- | --- |
| Diffusion weighted EPI | 33 | 5000 | 7/15 | 5 | 30 | 1000, 2500, 4000 | 0.1 x 0.1 x 0.7 | 2/15 min |

### 2.3 Image Analysis

MR images were reconstructed from raw data on the scanner console (Paravision 6.0.1, Bruker Biospin, Billerica, MA, USA). T_1_ and T_2_ values were computed by fitting the corresponding mono-exponential decay models to signals at each voxel using the Image Sequence Analysis (ISA) Tool in Bruker Paravision. Images with no saturation pulse (M0) were acquired with images with positive/negative or dual-frequency saturation pulses (*M*_+_, *M*_−_, and *M*_±_). MTR was calculated as: 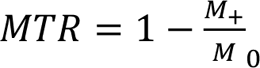 , and ihMTR was calculated as: 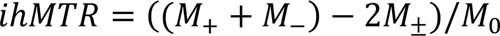. From diffusion MRI data, diffusion tensors and kurtosis tensors were obtained per voxel (Jensen et al., 2005) using the standard kurtosis-fitting approach implemented in the Diffusion Kurtosis Imaging Matlab Toolbox (https://cai2r.net/resources/software/diffusion-kurtosis-imaging-matlab-toolbox) (Veraart et al., 2013) and the following parameters were derived: mean and radial diffusivity (MD/RD), fractional anisotropy (FA), mean and radial kurtoses (MK/RK). The FA image was used for selecting the region of interest (ROI) using ROIeditor (http://www.mristudio.org) due to its superior contrast for white matter structures in the P10 mouse brains than conventional T_1_ and T_2_-weighted MRI. The ROIs included the genu and splenium of corpus callosum, the external capsule, cerebral peduncle, internal capsule for white matter (WM) structures and the motor cortex, sensory cortex, visual cortex, thalamus, caudate putamen, and hypothalamus for gray matter structures.

### 2.4. Histology

Post MRI *ex vivo* brains were dissected, cryo-protected in 30% sucrose solution and frozen embedded in OCT. Frozen brains were sectioned at 20 um thickness on a cryostat. Slides contain sections from the rostral to caudal regions of the corpus callosum. Slides were washed in PBS, incubated in 100% methanol at -20 °C for 10 minutes, incubated in blocking solution (10% goat serum diluted in 1X PBS, 1% BSA, 0.25% Triton-X100) for 1 hour at room temperature. Slides were incubated with the primary antibody (diluted in 1X PBS, 1% BSA, 0.25% Triton-X100) overnight at 4 °C. Slides were rinsed 3 x 5 minutes with PBS then incubated in the Alexa Fluor secondary antibody for 1 hour at room temperature. Slides were rinsed 3 x 5 minutes with PBS and a final wash in distilled water then mounted with Fluoromount G mounting media and cover slipped. The primary antibody used for immunohistochemistry was chicken anti-MBP (Millipore, AB9348, 1:200); the secondary antibody used was goat anti-chicken (IgG 488, Jackson ImmunoResearch, 1:1000). Slides were imaged and acquired at 20x on Confocal Zeiss LSM 800 Fluorescence Microscope using Zeiss Zen blue software. Using ImageJ (Schneider et al., 2012), ROIs were defined that matched the ROIs defined in MR.I

### 2.5. Statistics

All statistical tests were performed with Prism (GraphPad). Statistical significance was determined using the Student’s *t*-test with the threshold set at 0.05. Pearson correlation coefficients and p values were calculated between MR parameters and MBP signals. Corrections for multiple comparisons were performed using the Bonferroni method. All values in bar graphs indicate mean values and standard deviations.

We used the function *plsregress* in Matlab® 9.6.0 (R2019a) to compute variable importance in projection (VIP) score, regression coefficients, and the percent of variance explained in the response variable (PCTVAR). The function projected the independent variables (MR parameters) and response variables (MBP signals) into a latent space, in which linear least square regression was carried out. This approach reduces collinearity within the independent variables and their dimension without reducing the prediction power (Yoshida et al., 2017).

## 3. Results

### 3.1 Multi-parametric MRI and histology of Gli1^-/-^ and control mice

**Fig. 1** and supplementary **Fig. S1** show representative MBP-stained histology of *Gli1^-/-^* and *Gli1^+/-^* mouse brains at P10 and maps of nine MR parameters at comparable locations from the same animal. Overall, the white matter (WM) and gray matter (GM) contrasts in these parameter maps were lower than those observed in the adult mouse brain due to low myelin content in the P10 mouse brain (myelination begins at ∼P8 in the mouse brain). T_1_ and T_2_ maps showed that the mouse corpus callosum (cc) at P10 had longer T_1_ and T_2_ values than the neighboring cortex. The cc had lower MTR values but higher ihMTR values than the cortex. In comparison, diffusion MR parameters showed better WM/GM contrasts, with higher FA, higher MK/RK, and lower MD/RD in the cc than the cortex.

**Fig. 1.**
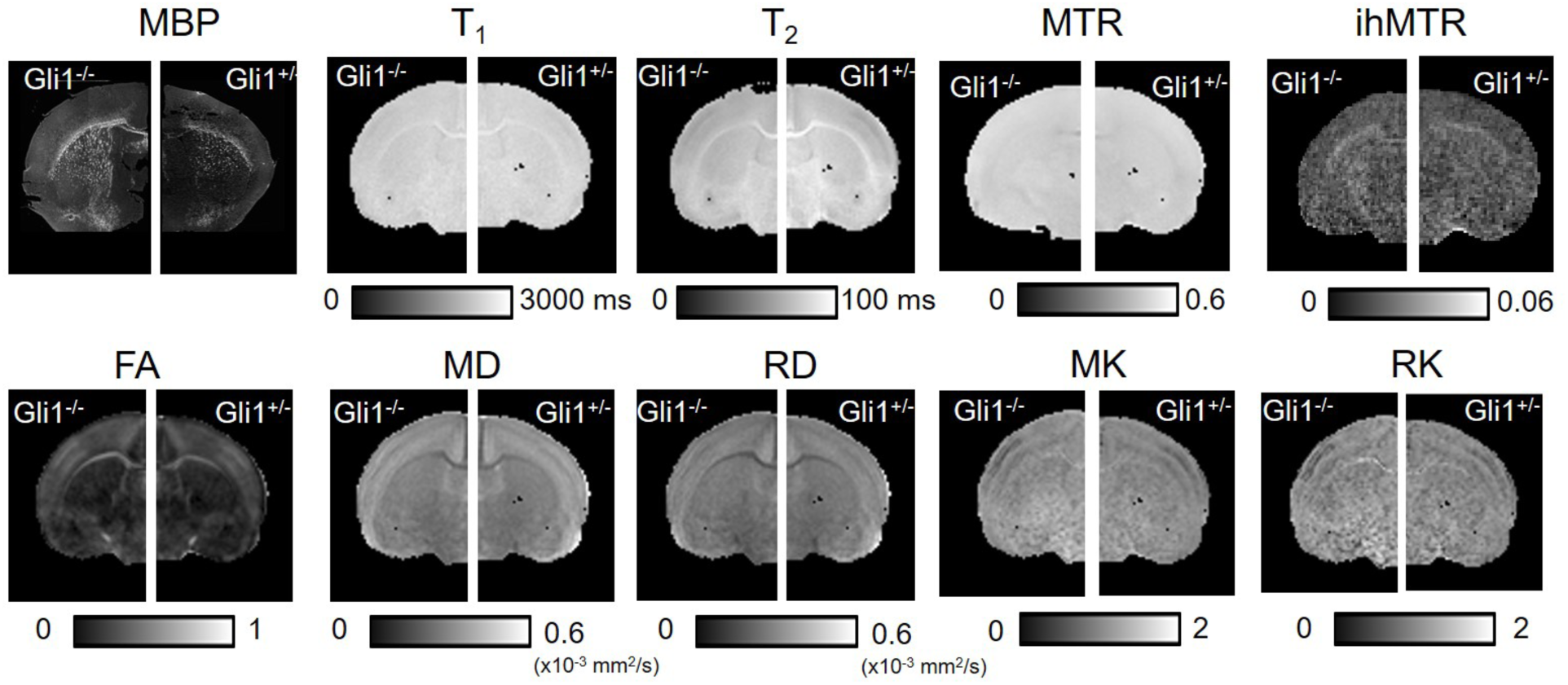
Representative axial MBP-stained histology of *Gli1^-/-^* and *Gli1^+/-^* mouse brains and corresponding MR parameter maps at the level of the genu of corpus callosum.

We first examined the correlations between MR parameters to mean intensity values in MBP-stained histology in manually defined ROIs. When we included data from multiple WM and GM ROIs in correlation analysis, we found that ihMTR, FA, and RK had moderate correlations with the MBP signals (*R* = 0.54, 0.57, and 0.54, respectively) (**Fig. 2A**). When only WM ROIs were considered (**Fig. 2B**), only MTR showed strong correlation with the MBP signals (*R* = 0.68). When only GM ROIs were considered (**Fig. 2C**), MTR and RD showed moderate correlations with the MBP signals (*R* = 0.50 and 0.54, respectively). Among MR parameters, we found strong correlation between MK and RK (R > 0.93), as well as between MD and RD (R >0.85) (**Fig. 2**).

**Fig. 2.**
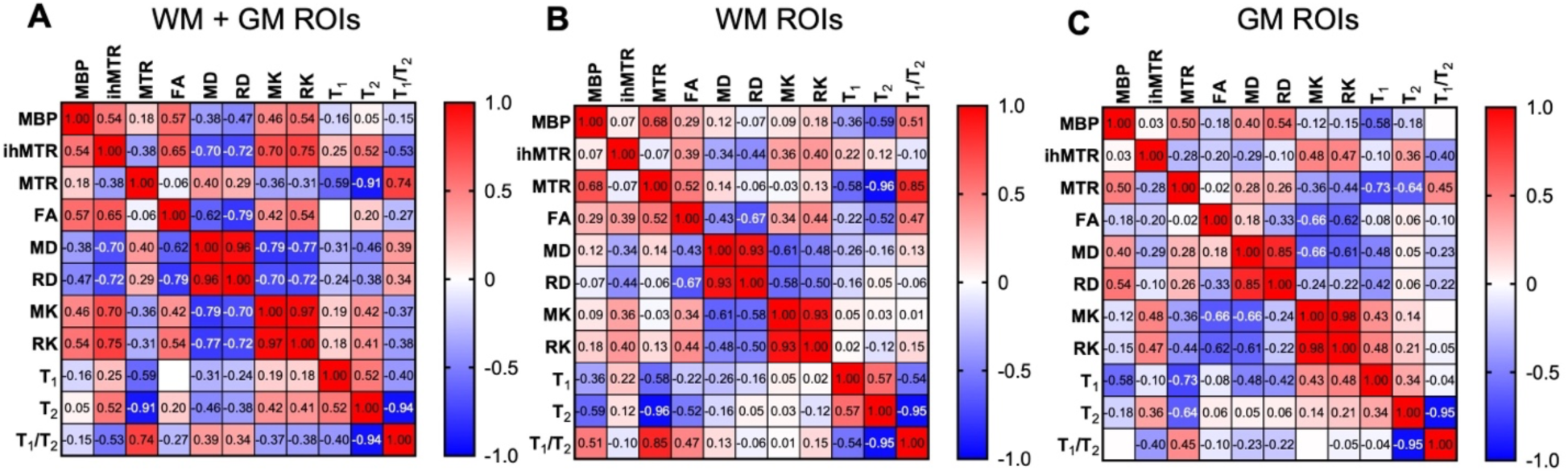
Correlation matrixes of multiple WM and GM ROIs in correlation analysis and averaged the results from both *Gli1^-/-^* and *Gli1^+/-^* mouse brains: WM and GM ROIs (A), WM ROIs (B) and GM ROIs (C). The values in the matrix are the correlation coefficients for significant correlations (*p*< 0.05) without corrections for multiple comparisons.

MBP values measured in both the genu and splenium of the corpus callosum (gcc and scc, respectively) confirmed enhanced myelination in the *Gli1^-/-^*mouse brains compared to the *Gli1^+/-^* mouse brains at P10 (**Fig. 3**). After corrections for multiple comparisons, most MR parameters showed significant differences between the *Gli1^-/-^* and *Gli1^+/-^* mouse brains in either the gcc or scc, but not in both.

**Fig. 3.**
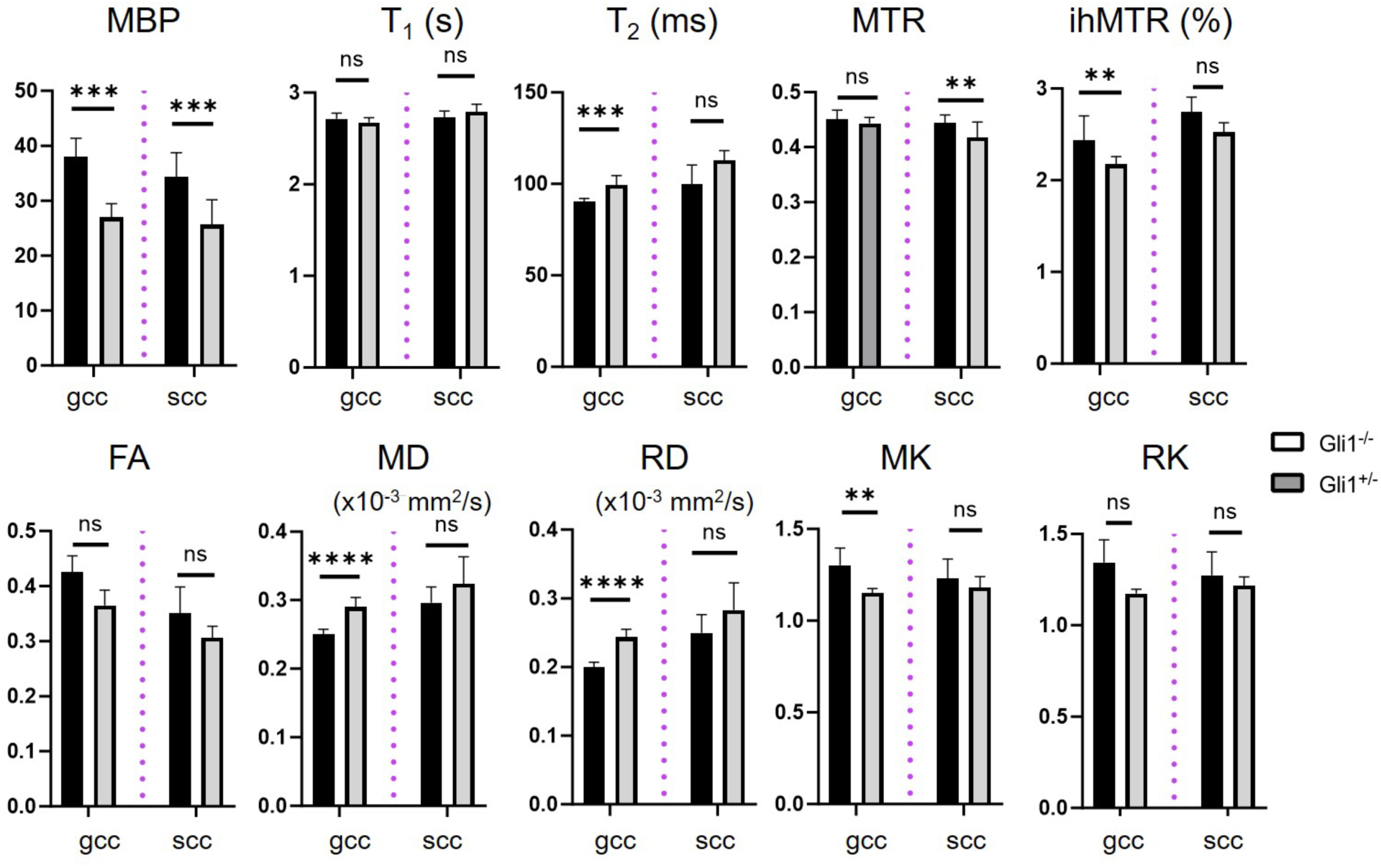
Quantitative analysis of MBP histology and MR parameter in gcc and scc in the *Gli1^-/-^* and *Gli1^+/-^* mouse brains. ***/****: *p* < 0.001/0.0001 by Student’s t-test without corrections for multiple comparisons. The threshold for significance was set at 0.005 to correct for multiple comparisons.

We calculated Pearson correlations between MBP signals and individual MR parameters in the corpus callosum. In the gcc (**Fig. 4**), RD had the strongest correlation with MBP (*R^2^=0.78, p=0.004*). T_1_/T_2_ and T_2_ showed relatively strong correlations with MBP (*R^2^=0.75, p=0.001*, and *R^2^=0.67, p=0.004*, respectively). Here, we used T_1_/T_2_ to include the interaction between these two parameters instead of the ratio between T_1_-and T_2_-weighted signals. MTR, ihMTR, FA, and RK had moderate correlations with MBP (*R^2^=0.52, p=0.02; R^2^=0.52, p=0.02; R^2^=0.55, p=0.01;* and *R^2^=0.50, p=0.02*, respectively). In addition, T_1_ had a weak correlation (*R^2^=0.19, p=0.22*). After corrections for multiple comparisons, only RD, T_2_, T_1_/T_2_ had significant correlations with MBP signals.

**Fig. 4.**
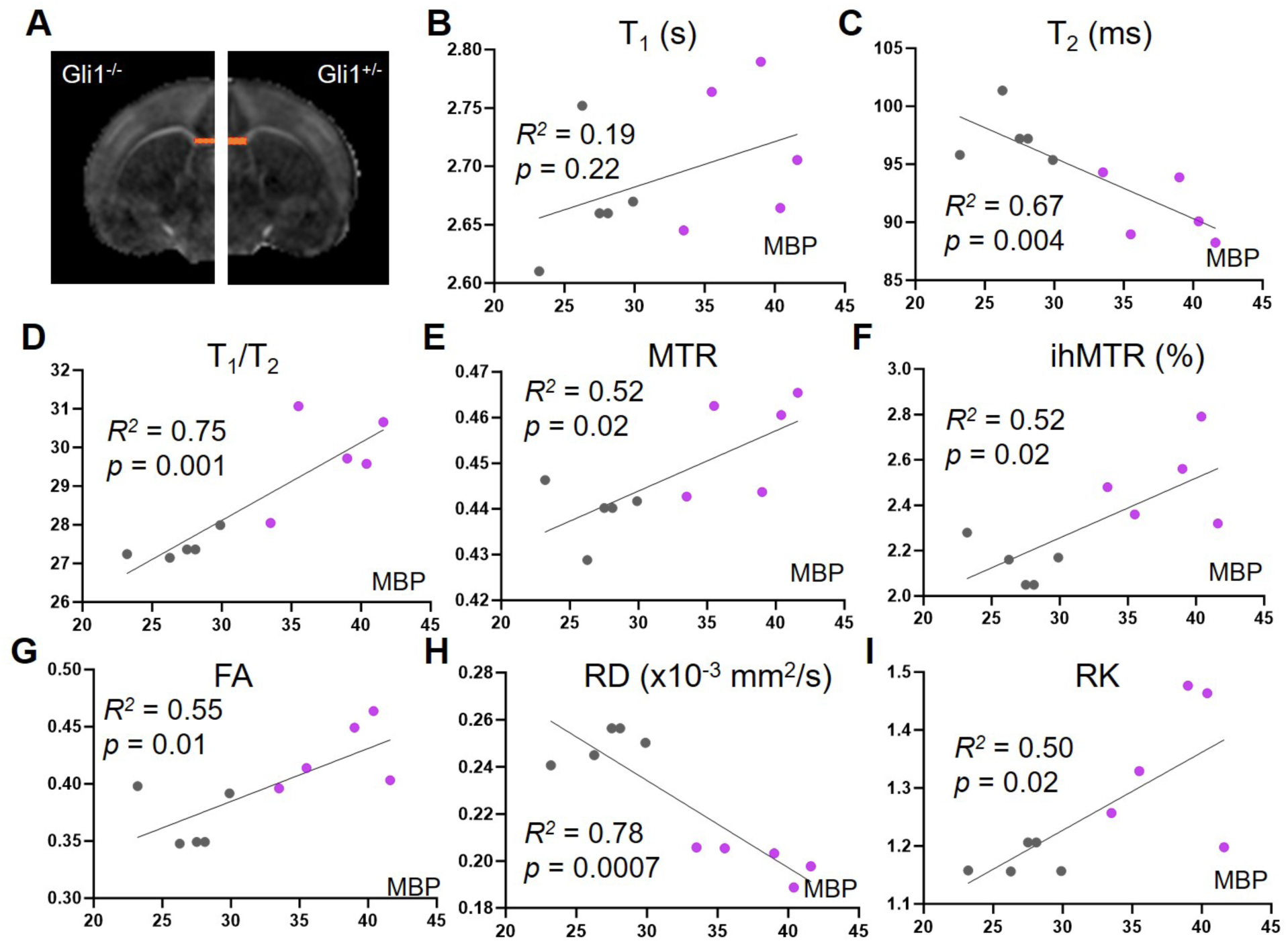
Correlations between MR parameters and MBP signals in the genu of the corpus callosum (gcc). **A:** An example of the ROI for gcc defined in the FA images. **B-I:** Scatter plots of MR parameters and MBP signals in the gcc of both *Gli1^-/-^* (purple dots) and *Gli1^+/-^* (gray dots) mouse brains. The *p* values shown here are before corrections for multiple comparisons.

### 3.2 Estimation of myelin in the gcc based on multiple MR parameters

For PLSR analysis, we used mean MR parameter values in the gcc from *Gli1^-/-^* and *Gli1^+/-^* mouse brain as independent variables and the corresponding MBP signals as response variables. Based on the results in **Figs. 2 and 4**, we only included ihMTR, FA, RD, RK, T_2_, T_1_/T_2_ to minimize redundancy and avoid over-fitting. The VIP scores suggested that T_2_ and T_1_/T_2_ had high contributions to myelin predictions (VIP score > 1), followed by ihMTR, whose VIP score was slightly lower than one (**Fig. 5A**). At least 3 latent components were needed to explain approximately 80% variance of the MBP signals (**Fig. 5B**), supporting the inclusion of multiple MRI parameters. When only T_2_, T_1_/T_2_, ihMTR were included for the PLSR analysis, a similar percentage in the variance of the MBP signals could be explained (**Fig. 5C**). The regression model from PSLR was *MBP = 26.49 + 2.73*ihMTR -0.19*T_2_ + 0.71*T_1_/T_2_*. The correlation between the estimated MBP based on the three MR parameters and the actual MBP signals was stronger (*R^2^=0.83, p=0.0002*, **Fig. 5D**) than any individual MR parameters (**Fig. 4**).

**Fig. 5.**
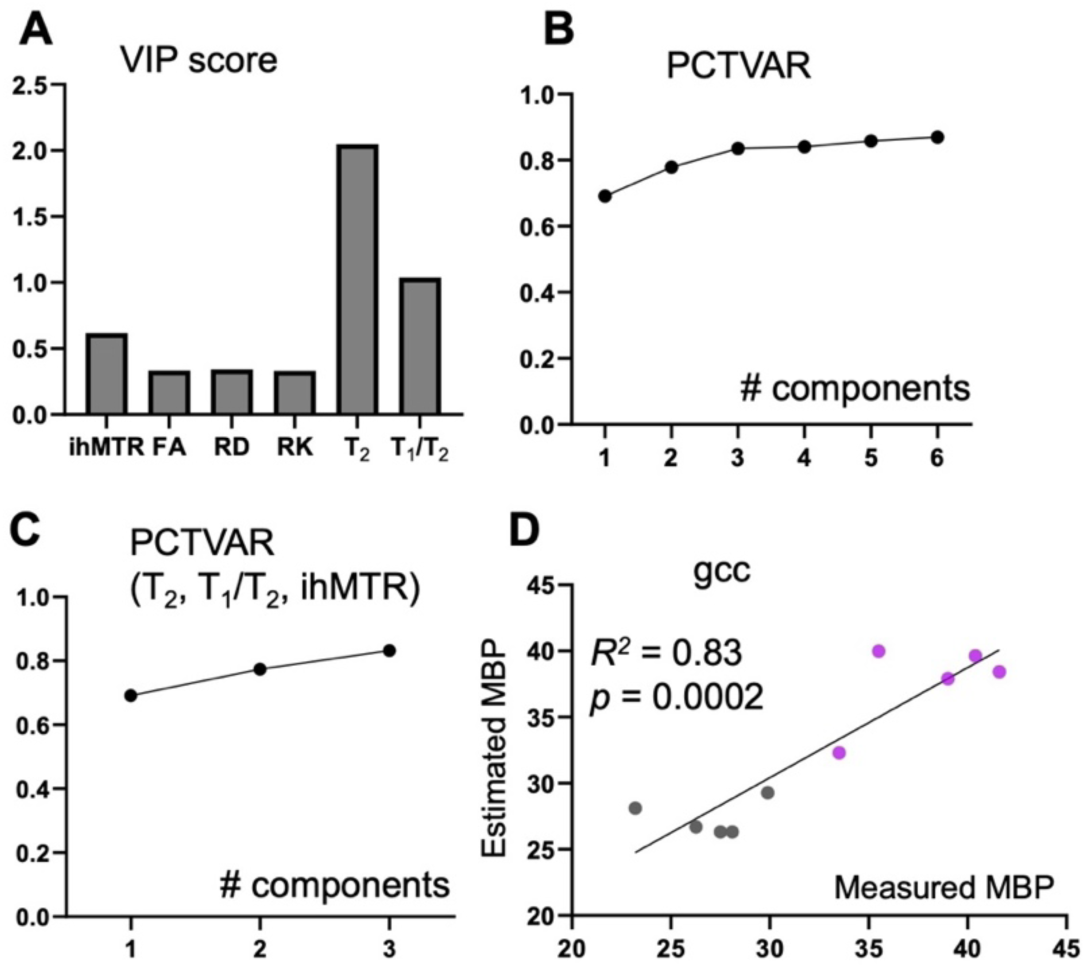
Estimation of myelin based on multiple MR parameters in the genu of corpus callosum (gcc). **A-B:** The variable importance in projection (VIP) scores of each MR parameter and the percentage of variance (PCTVAR) of the MBP signals in the gcc that can be explained by latent components. **C:** The PCTVAR of the MBP signals in the gcc that can be explained by three latent components from T_2_, T_1_/T_2_, and ihMTR. **D:** The scatter plot of estimated MBP signals based on T_2_, T_1_/T_2_, ihMTR and measured MBP signals in the gcc from *Gli1*^-/-^ (purple dots) and *Gli1*^+/-^ (gray dots) mouse brains.

### 3.3. Estimation of myelin in the motor cortex based on multiple MR parameters

In the motor cortex (mCX), T_2_ had a strong correlation with MBP (*R^2^=0.66, p=0.01*), with MTR and T_1_/T_2_ showed moderate correlations with MBP (*R^2^=0.56, p=0.01*, and *R^2^=0.55, p=0.02*, respectively) (**Fig. 6**). However, after corrections for multiple comparisons, none of the MR parameters was significantly correlated with the MBP signals. The VIP scores from PLSR suggested that T_2_ and RK had the highest contribution to myelin prediction (VIP score > 1), followed by T_1_/T_2_, whose VIP score was slightly lower than one (**Fig. 7A**). At least two latent components were needed to explain approximately 70% variance of the MBP signals, and three to reach more than 90% (**Fig.7B**). When only T_2_, RK, and T_1_/T_2_ were included, more than 90% variance in the MBP signals could be explained (**Fig. 7C**). The regression model from PLSR for MBP in the motor cortex was *MBP = 182.33 + 53.64*RK - 1.89*T_2_ -2.2*T_1_/T_2_*. Based on the three MR parameters, the estimated MBP showed a strong and significant correlation with the actual MBP signals (*R^2^=0.95, p<0.0001*, **Fig. 7D**).

**Fig. 6.**
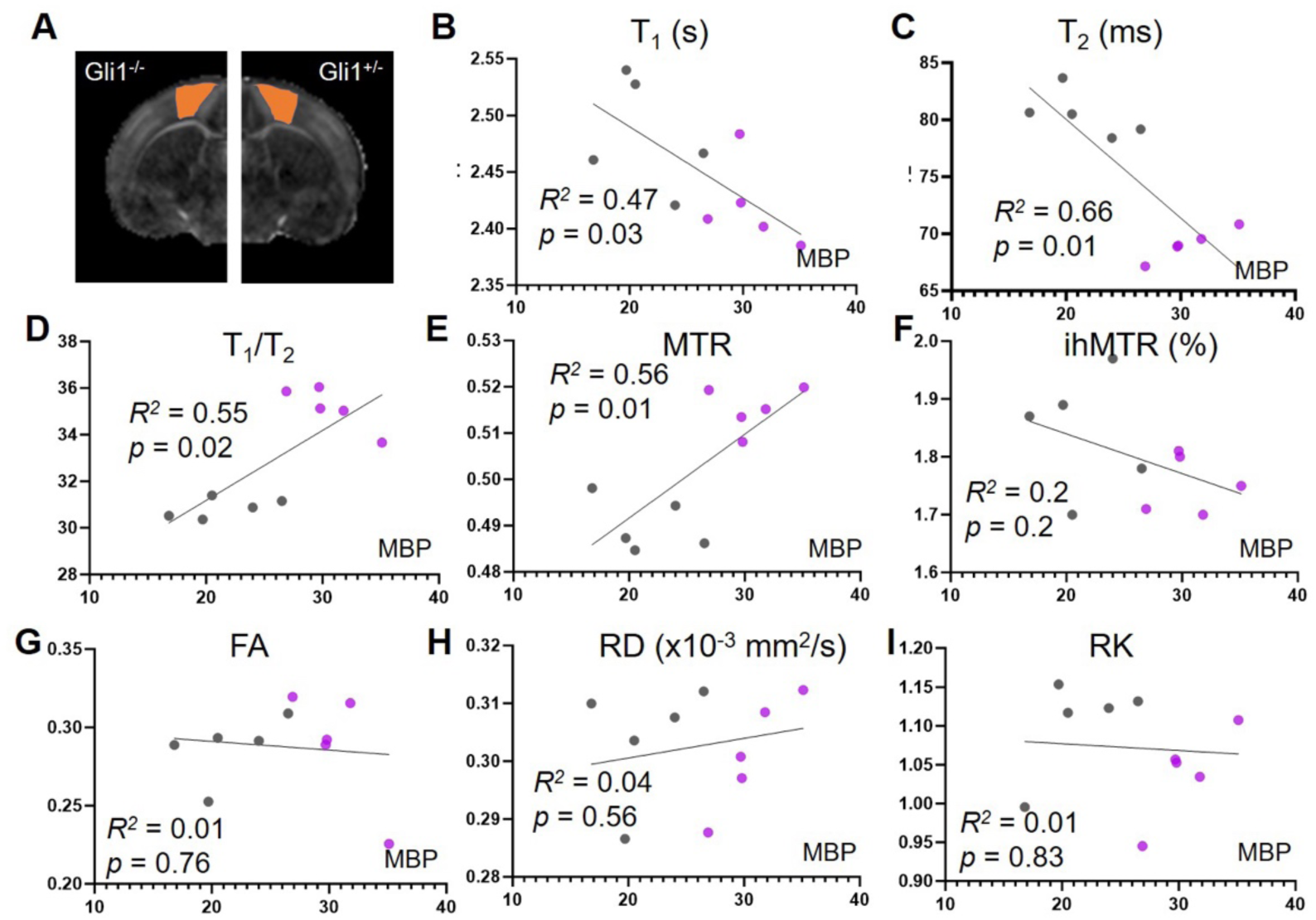
Correlations between MR parameters and MBP signals in the motor cortex (mCX). **A:** An example of the ROI for mCX defined in the FA images. **B-I:** Scatter plots of MR parameters and MBP signals in the gcc of both *Gli1^-/-^* (purple dots) and *Gli1^+/-^* (gray dots) mouse brains.

**Fig. 7.**
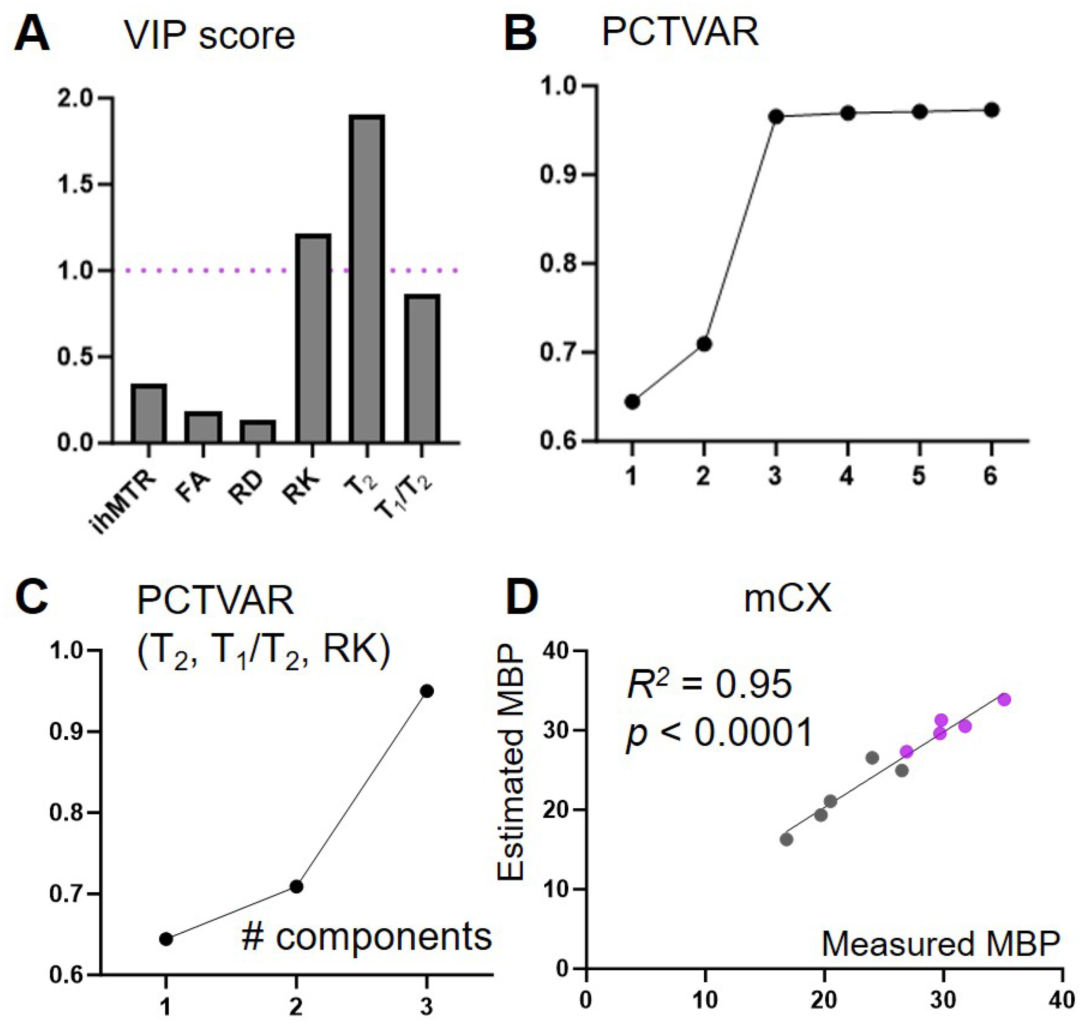
Estimation of myelin based on multiple MR parameters in the motor cortex (mCX). **A-B:** The variable importance in projection (VIP) scores of each MR parameter and the percentage of variance (PCTVAR) of the MBP signals in the mCX that can be explained by the number of latent components. **C:** The PCTVAR of the MBP signals in the mCX that can be explained by three latent components from T_2_, T_1_/T_2_, and RK. **D:** The scatter plot of estimated MBP signals based on T_2_, T_1_/T_2_, RK and measured MBP signals in the mCX from *Gli1*^-/-^ (purple dots) and control (gray dots) mouse brains.

## 4 Discussion

In this study, we explored the use of multiparametric MRI for measuring myelin changes in the *Gli1^-/-^* mice, a model of accelerated myelination. Our findings indicate that multi-parametric MRI can improve the precision of myelin mapping. Specifically, we employed PLSR to evaluate and rank MR parameters based on their contributions to myelin estimation, thereby facilitating the refinement of multi-parametric MRI based myelin mapping. Furthermore, our study suggests that the mapping of myelin across brain regions with different microstructural organization (e.g., levels of myelin and axon) may require distinct MR parameter sets, underscoring the challenges in developing a universal myelin marker applicable to the entire brain.

### 4.1 Correlations between MR parameters and MBP histology in the *Gli1* mouse model

Although correlations have been frequently used to examine the relationship between MR parameters and myelin, previous studies have demonstrated that the observed correlations might be driven more by the diverse regional microstructural organizations than variations in myelin content and its structural integrity, when multiple brain regions within the subjects are included (Lazari and Lipp, 2021; Mancini et al., 2020). Results in **Fig. 2** demonstrate that the correlations between MR parameters and MBP signals indeed depend on the regions examined. To reduce the potential influence of regional microstructural organization, this study focused on the corpus callosum and the cortex separately. The corpus callosum is comprised primarily of coherently arranged axons, whereas a large portion of the cortex is occupied by soma and neuropils, with less coherent organizations. The corpus callosum and cortex, therefore, represent two distinct cases. Furthermore, the *Gli1* mouse model employed here, unlike mouse models commonly studied by MRI for detecting myelin injury and repair (e.g., the cuprizone model (Matsushima and Morell, 2001) or EAE (Constantinescu et al., 2011), does not exhibit inflammation or axonal damage, thereby further limiting the impacts of non-myelin related factors. However, it is important to note that complete removal of such influences, for instance, variations in axonal density in the corpus callosum, remains challenging. This may account for the observation that, while most MR parameters included in this study showed significant differences in the corpus callosum between the *Gli1^-/-^*and *Gli1^+/-^* mouse brains (**Fig. 3**), only a subset of them (T_2_, T_1_/T_2_, and RD) demonstrated significant correlations with MBP-stained histology (**Fig. 4**), as some of the observed differences could indicate potential variations in axonal densities or other aspects of tissue microstructure.

### 4.2 Does a universal MRI-based myelin marker exist?

Early studies had introduced the use of multi-parametric MRI to enhance myelin mapping in the brain (Mangeat et al., 2015; Patel et al., 2020; Stikov et al., 2015; Stikov et al., 2011). This premise is further supported by recent deep learning-based studies. One study reported a deep learning network that can predict myelin water fraction, a metric with greater myelin specificity than conventional MR parameters, based on conventional T_1_, T_2_, and diffusion MR parameters (Drenthen et al., 2021). We have shown that a deep learning network, trained on co-registered multi-parametric MRI data (including T_2_-weighted, MTR, and diffusion MR parameters) and co-registered MBP-stained histology can improve myelin specificity (Liang et al., 2022). Despite these advancements, the inner-working of these networks remain largely elusive.

Considering the value of multi-parametric MRI in myelin mapping, it is important to examine the contributions of individual MR parameters for further optimization, e.g., to pick the most useful ones while discarding those that are redundant can shorten acquisition time, as gathering multi-parametric MRI data can be time-consuming. In this study, we employed PLSR, a method capable of handling collinearity among MR parameters, to evaluate the contributions of MR parameters and derive a linear myelin estimator based on a subset of MR parameters with high contributions. The estimator is basically a local linear approximation of the complex relationship between MR parameters and myelin, along with its surrounding microstructural organization. In the corpus callosum, T_2_ and T_1_/T_2_ were the only parameters with VIP score higher than 1, followed by ihMTR. This is not surprising, as T_2_ and T_1_/T_2_ signal showed strong correlation with MBP signals in the corpus callosum (Fig. 4). Previous reports have suggested that relaxivity and MT parameters are more robust myelin predictors than diffusion parameters (Lazari and Lipp, 2021; Mancini et al., 2020). The relatively low contribution from MT parameters could be caused by the limited myelin in the P10 mouse brain, especially macromolecules and lipids rich in fully myelinated white matter, as evidenced by the weak contrasts in MTR and ihMTR maps (**Fig. 1**).

When we applied the same approach to the data from the motor cortex, the results differed slightly from those observed in the corpus callosum. T_2_ again had high contributions (VIP scores > 1), yet RK emerged as more important than either MTR or ihMTR. Given the lower myelin content and unique microstructural organization of the motor cortex relative to the corpus callosum, it is plausible that the linear approximation of the relationship between myelin and MR parameters is altered as well. In the motor cortex, RK may reflect the density of axons in the motor cortex, providing valuable insights for myelin estimation that might not be as critical in the corpus callosum. These findings underscore the difficulty in identifying a universal MRI-based marker for myelin. Due to the diverse microstructural organizations across different brain areas—and even within the same region across various stages of development and disease—the relationship between myelin content and MRI parameters is subject to change. This necessitates comprehensive studies encompassing both healthy and diseased states to better understand these relationships.

### 4.3. Limitations of the current study

This study has several limitations. Both the choice of *ex vivo* MRI, which is known to differ from *in vivo* MRI (Roebroeck et al., 2019; Shepherd et al., 2009; Zhang et al., 2012), and the early stage of myelination at P10, make it difficult to generalize the findings here to *in vivo* studies and mouse brains at later stages. This is necessary because the enhanced myelination observed in the *Gli1^-/-^* mouse brain at P10 may not sustain to later stages and *in vivo* multi-parametric MRI of the P10 mouse is challenging. In addition, there are other key aspects of myelin (e.g., myelin lipids) not captured by the MBP-stained histology, and variations due to the histology preparation have not been examined here. It is reasonable to expect certain variances in MBP signals coming from tissue processing (including fixation and dehydration), and staining, which may be reduced in future studies by utilizing recent advances in tissue clearing (Richardson et al., 2021) and genetically modified mouse models with GFP co-expressed with selected myelin proteins (Hughes et al., 2018; Mallon et al., 2002). As obtaining myelin histology for a large number of samples is time consuming, we only included a limited number of animals in this study, which precluded testing the findings here in a separate cohort of animals. Future studies including a larger cohort, animals at later stages of myelination, and other immunohistochemical markers for myelin will be useful to further enhance our understanding in this area.

Even though we included a relatively large set of MRI parameters in this study, several important MRI-based myelin markers were left out due to practical reasons. One such marker is myelin water fraction from myelin water imaging, which has been shown to be a robust marker of remyelination in a clinical trial (Caverzasi et al., 2023). In the P10 mouse brain, it is unknown whether compact myelin has formed at this stage and what the myelin water T_2_ values are, making it challenging to estimate the fraction of myelin water compartment and will require further validation and optimization. Quantitative Magnetization Transfer (qMT) is another marker that was left out. qMT has demonstrated its ability to effectively assess demyelination through the pool size ratio in a shiverer mouse model (Ou et al., 2009) and has shown strong correlations with histopathological findings in a rabbit model (Drobyshevsky et al., 2023). Incorporating these advanced myelin markers in future studies will be a high priority.

## 5 Conclusion

Our results demonstrated that multi-parametric MRI can detect enhanced myelination in the *Gli1* mouse model. Based on MBP-stained myelin histology, the relationship between MR parameters and myelin was studied in the corpus callosum and motor cortex, and the contributions of MR parameters were evaluated.

## Data and Code Availability

The following openly available software were used:

Diffusion Kurtosis Imaging Matlab Toolbox (https://cai2r.net/resources/software/diffusion-kurtosis-imaging-matlab-toolbox)

ROIeditor (http://www.mristudio.org)

Prism (GraphPad) (https://www.graphpad.com)

## Author Contributions

C.H.L. conceived the study, collected the data, performed all analyses, and wrote the initial draft of the manuscript. M.H performed histology. J. L.S. conceived the study, contributed to study design and interpretation of histology data. J.Z. contributed to the hypothesis generation and study design, supervised and participated in data collection and analysis. All authors reviewed and edited the manuscript.

## Acknowledgement

We thank Jacob Brady for assisting in sample preparation and histology. This study was supported by the National Institute of Health R01NS102904, R01HD074593, U24NS135568 to (JZ) and R01NS100867 (JLS). MR imaging was performed at the Center of Advanced Imaging Innovation and Research (CAI2R, www.cai2r.net), a Biomedical Technology Resource Center supported by NIBIB with the award P41 EB017183.

## Notes

### Competing Interest Statement

The authors have declared no competing interest.

### Summary of Updates

The results have been updated and extended, and the discussion section has also been revised.

