## Supplemental figures for "Multi-parametric MRI can detect enhanced myelination in the *Gli1^-/-^* mouse brain"

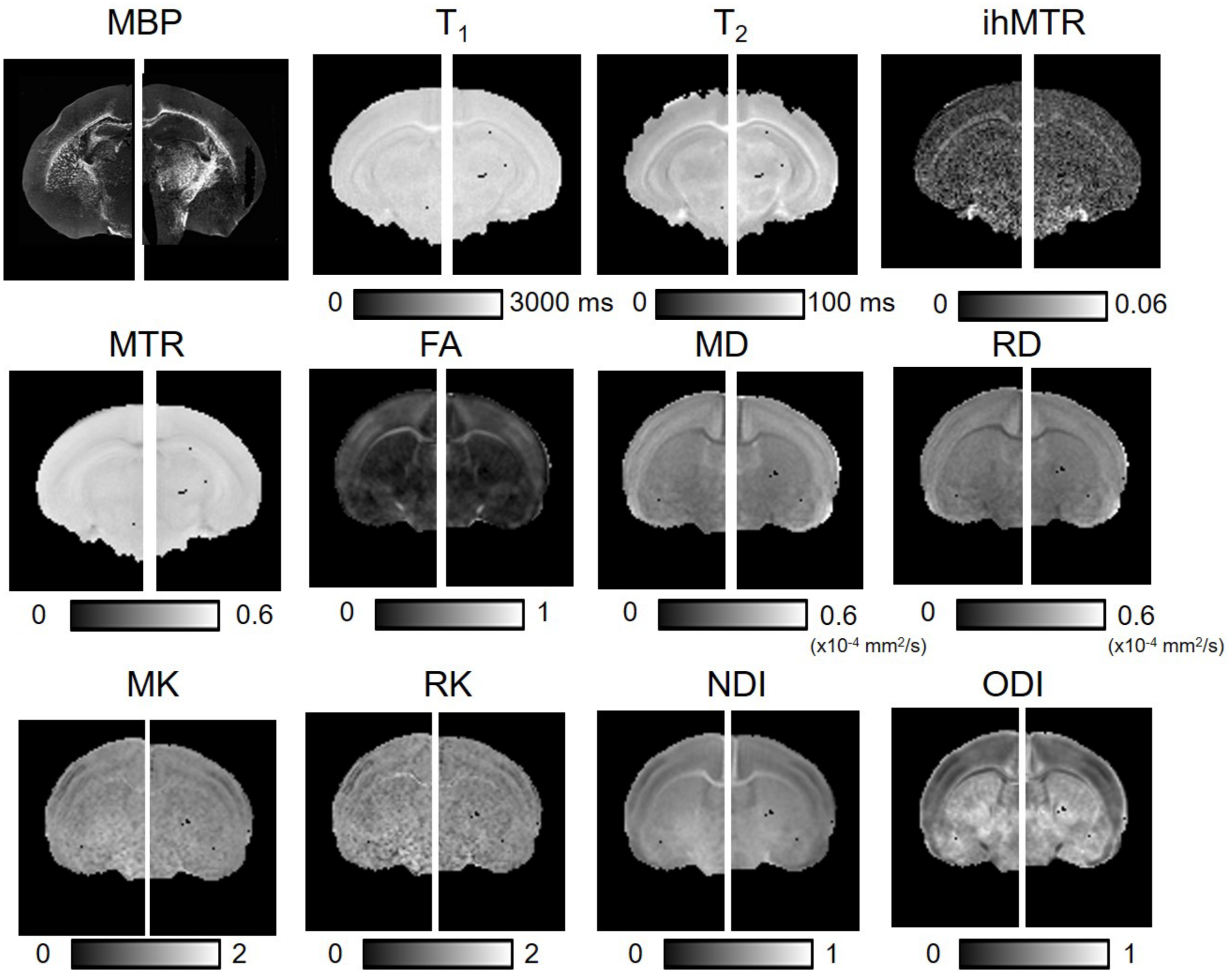


**Fig.S1:** Representative axial MBP-stained histology of *Gli1*$^{-/-}$ and *Gli1*$^{+/-}$ mouse brains and corresponding MR parameter maps at the level of the splenium of corpus callosum.

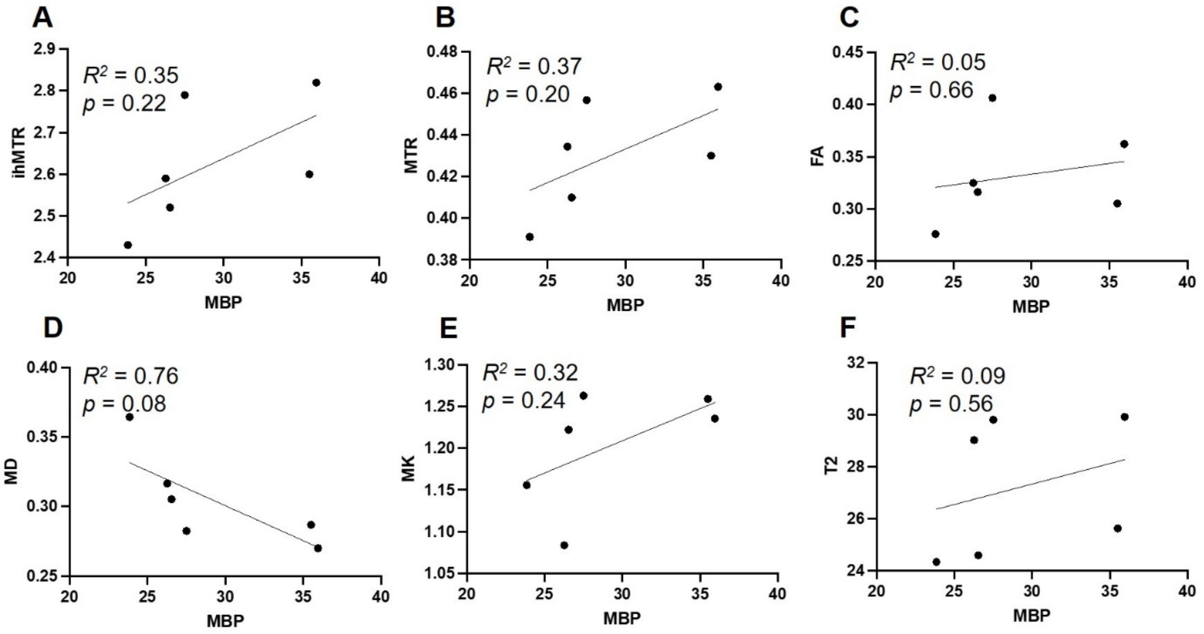


**Fig.S2:** Correlation between MBP signals and MR parameters in the splenium of corpus callosum (scc) of *Gli1*$^{-/-}$ and *Gli1*$^{-/+}$ mouse brains.
